# A new class of receptors: Lipids regulate mammalian Gsα-stimulated adenylyl cyclase activities via their membrane anchors

**DOI:** 10.1101/2024.08.06.606792

**Authors:** Marius Landau, Sherif Elsabbagh, Harald Gross, Adrian Fuchs, Anita C.F. Schultz, Joachim E. Schultz

**Author notes:** Corresponding author: Dr. Joachim E Schultz Pharmazeutisches Institut der Universität Tübingen, Auf der Morgenstelle 8, 72076 Tübingen, Germany.

## Abstract

The biosynthesis of cAMP by mammalian membrane-bound adenylyl cyclases (mACs) is predominantly regulated by G-protein-coupled-receptors (GPCRs). Up to now the two hexahelical transmembrane domains of mACs were considered to fix the enzyme to membranes. Here we show that the transmembrane domains serve in addition as signal receptors and transmitters of lipid signals that control Gsα-stimulated mAC activities. We identify aliphatic fatty acids and anandamide as receptor ligands of mAC isoforms 1 to 7 and 9. The ligands enhance (mAC isoforms 2, 3, 7, and 9) or attenuate (isoforms 1, 4, 5, and 6) Gsα-stimulated mAC activities *in vitro* and *in vivo*. Substitution of the stimulatory membrane receptor of mAC3 by the inhibitory receptor of mAC5 results in a ligand inhibited mAC5-mAC3 chimera. Thus, we discovered a new class of membrane receptors in which two signaling modalities are at a crossing, direct tonic lipid and indirect phasic GPCR-Gsα signaling regulating the biosynthesis of cAMP.

---

The structure of the first second messenger, cyclic 3‘,5‘-adenosine monophosphate (cAMP), was reported in 1958 by Sutherland and Rall. It was generated by incubation of a liver extract with the first messengers adrenaline or glucagon (1). Since then, cAMP has been demonstrated to be an almost universal second messenger used to translate various extracellular stimuli into a uniform intracellular chemical signal (2, 3). The biosynthetic enzymes for cAMP from ATP, adenylyl cyclases (ACs), have been biochemically investigated in bacteria and eukaryotic cells (4–6). Finally in 1989, the first mammalian AC was sequenced (7). The protein displayed two catalytic domains (C1 and C2) which are highly similar in all isoforms. The two hexahelical membrane anchors (TM1 and TM2) are dissimilar and isoform specifically conserved (8). The latter were proposed to possess a channel or transporter-like function, properties, which were never confirmed (7). The domain architecture of mammalian ACs, TM1-C1-TM2-C2, clearly indicated a pseudoheterodimeric protein composed of two concatenated monomeric bacterial precursor proteins (9). To date, sequencing has identified nine mammalian membrane-delimited AC isoforms (mACs) with identical domain architectures (2). In all nine isoforms the catalytic domains are conserved and share extensive sequence and structural similarities which indicate a similar biosynthetic mechanism (5, 6, 10, 11). On the other hand, the associated two membrane domains differ substantially within each isoform and between all nine mAC isoforms (8, 12, 13). Bioinformatic studies, however, revealed that the membrane domains, TM1 as well as TM2, are highly conserved in an isoform-specific manner for about 0.5 billion years of evolution (8, 12).

Extensive studies on the regulation of the nine mAC isoforms revealed that the Gsα subunit of the trimeric G-proteins activate cAMP formation. Gsα is released intracellularly upon stimulation of G-protein-coupled receptors (GPCR), i.e. the receptor function for mAC regulation was assigned to the diversity of GPCRs which presently are most prominent drug targets. In 1995 Tang and Gilman reported that Gsα regulation of mammalian mACs does not require the presence of the membrane anchors. A soluble C1-C2 dimer devoid of the membrane domains was fully activated by Gsα (13). This reinforced the view that the mAC membrane anchors which comprise up to 40 % of the protein were just that and otherwise functionally inert. The theoretical possibility of mACs to be regulated directly, bypassing the GPCRs, was dismissed (14).

We were intrigued by the exceptional evolutionary conservation of the mAC membrane anchors for 0.5 billion years (8, 12). In addition, we identified a cyclase-transducing-element that connects the TM1 and TM2 domains to the attached C1 or C2 catalytic domains. These transducer elements are similarly conserved in a strictly isoform-specific manner (12, 15). Furthermore, cryo-EM structures of mAC holoenzymes clearly revealed that the two membrane domains, TM1 and TM2, collapse into a tight dodecahelical complex resembling membrane receptors (16–18). Lastly, we created a chimeric model involving the hexahelical quorum-sensing receptor from *Vibrio cholerae* which has a known aliphatic lipid ligand and the mAC 2 isoform. We observed that the ligand directly affected, i.e., attenuated the Gsα activation of mAC2 (19). As a proof-of concept this demonstrated that the cytosolic catalytic AC dimer serves as a receiver for extracellular signals transmitted through the dyad-related membrane anchor. Accordingly, we proposed a general model of mAC regulation in which the extent of the indirect mAC activation via the GPCR/Gsα axis is under direct ligand control via the mAC membrane anchor (19).

Here we report the results of a rigorous search for potential ligands of the mAC membrane domains. We identified aliphatic lipids as ligands for mAC isoforms 1 to 7 and 9. Isoform-dependently, the ligands either attenuate or enhance Gsα-activation *in vitro* and *in vivo* demonstrating a receptor function of the mAC proteins. The receptor properties are transferable as demonstrated by interchanging the membrane anchors between mAC3 and 5. Thus, the results define a new class of membrane receptors and establish a completely new level of regulation of cAMP biosynthesis in mammals in which tonic and phasic signaling processes intersect in a central signaling system, which is the target of frequently used drugs.

## Oleic acid enhances Gsα-stimulated mAC3, but not mAC5 activity

In earlier experiments we demonstrated regulation of the mycobacterial AC Rv2212 by lipids (20), the presence of oleic acid in the mycobacterial AC Rv1264 structure (21), and regulation of a chimera consisting of the quorum-sensing receptor from *Vibrio* and the mAC2 catalytic dimer by the aliphatic lipid 3-hydroxytridecan-4-one (22). Therefore, we searched for lipids as ligands. Here, we used bovine lung as a starting material because lipids are important for lung development and function (23, 24). Lipids were extracted from a cleared lung homogenate, acidified to pH 1, with dichloromethane/methanol (2:1). The dried organic phase was chromatographed on silica gel (employing vacuum-liquid-chromatography; Si-VLC) and fractions A to Q were assayed (fractionation scheme in appendix 1). Fraction E enhanced mAC3 activity stimulated by 300 nM Gsα four-fold, whereas mAC5 activity was unaffected (Fig.1A). The non-maximal concentration of 300 nM Gsα was used because it enabled us to observe stimulatory as well as inhibitory effects.

**Fig. 1.**
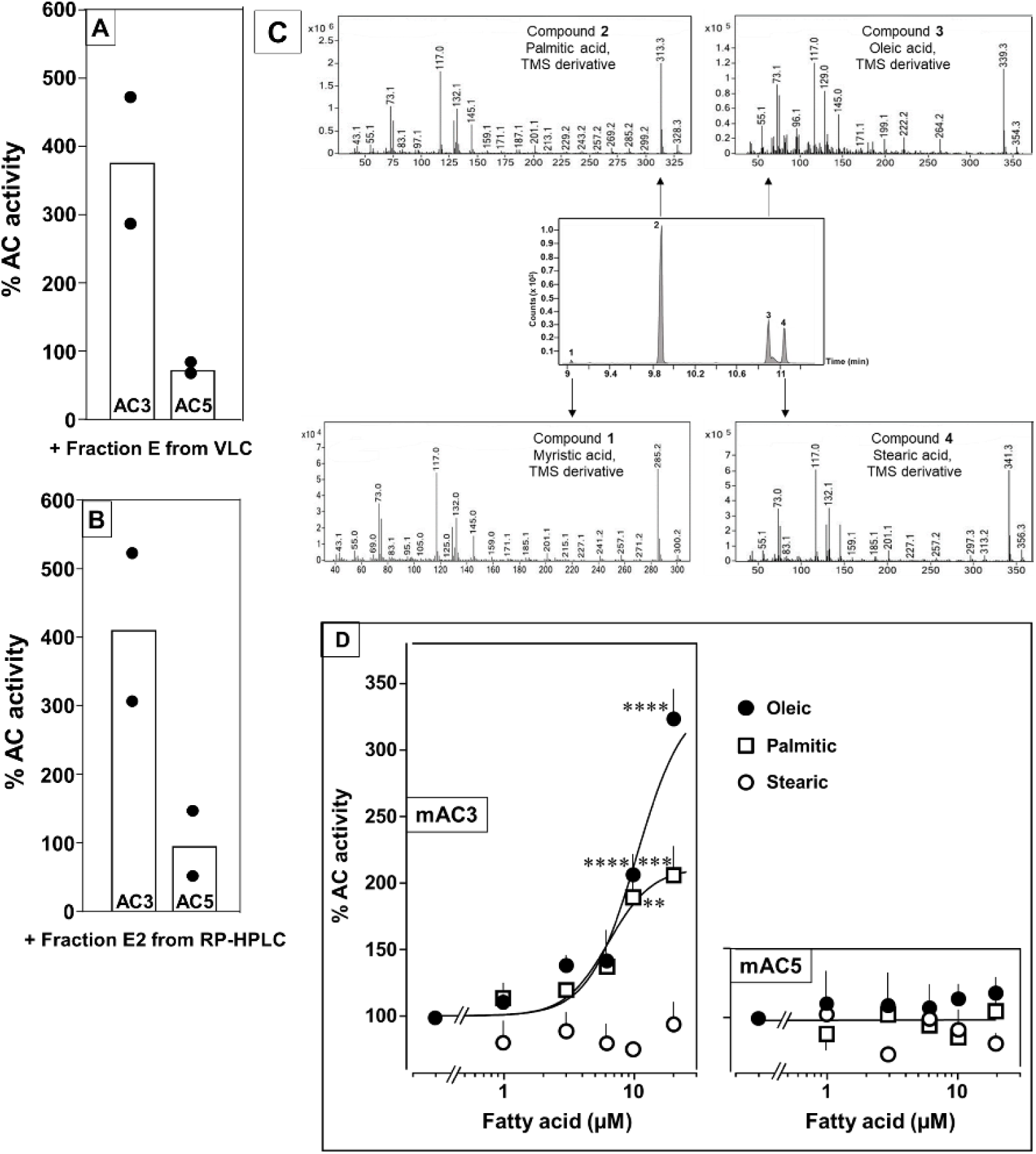
Identification of mAC3 activating fatty acids. **(A** and **B)** Effect of 1 µg/assay of fractions E from vacuum-liquid-chromatography (A) and E2 from RP-HPLC (B) on 300 nM Gsα-stimulated mAC isoforms 3 and 5. Activities are shown as % compared to 300 nM Gsα stimulation (100%). n=2, each with two technical replicates. Basal and Gsα activities of mAC3 in (A) were 0.01 and 0.07 and of mAC5 0.06 and 1.32 nmol cAMP•mg^-1^•min^-1^, respectively. In (B), basal and Gsα activities of mAC3 were 0.02 and 0.12 and of mAC5 0.09 and 1.1 nmol cAMP•mg^-1^•min^-1^, respectively. **(C)** GC-MS chromatogram of fraction E2. Mass spectra of the fatty acids are shown. Fatty acids’ identity was confirmed by comparing with corresponding standards (TMS: Trimethylsilyl). **(D)** Effect of fatty acids identified by GC-MS on 300 nM Gsα-stimulated mAC3 (left) and mAC5 (right). Basal and Gsα activities of mAC3 were 0.023 ± 0.02 and 0.17 ± 0.03 and of mAC5 0.08 ± 0.02 and 0.44 ± 0.09 nmol cAMP•mg^-1^•min^-1^, respectively. n= 3-23. EC_50_ of palmitic and oleic acids for mAC3 were 6.4 and 10.4 μM, respectively. Data represent individual experiments (black dots in A and B) or mean ± SEM (D). One-sample *t* tests were performed. Significances: \*\**P* < 0.01, \*\*\**P* < 0.001; \*\*\*\**P* < 0.0001.

Fraction E was further separated by RP-HPLC into five subfractions (E1 – E5; appendix 1 Fig. 1). The mAC3 enhancing constituents appeared in fraction E2. It enhanced Gsα-stimulated mAC3 four-fold but had no effect on mAC5 (Fig. 1B). ^1^H and ^13^C-NMR spectra of fraction E2 indicated the presence of aliphatic lipids (appendix 1 Fig. 2). Subsequent GC/MS analysis identified palmitic, stearic, oleic and myristic acid in E2 (Fig. 1C). Concentration-response curves were established for these fatty acids with mAC3 and mAC5 stimulated by 300 nM Gsα (Fig. 1D). 20 µM oleic acid enhanced Gsα-stimulated mAC3 activity three-fold (EC_50_ = 10.4 µM) and 20 µM palmitic acid two-fold (EC_50_ = 6.4 µM), while stearic or myristic acid had no significant effect. None of these fatty acids affected mAC5 activity (Fig. 1D).

**Fig. 2.**
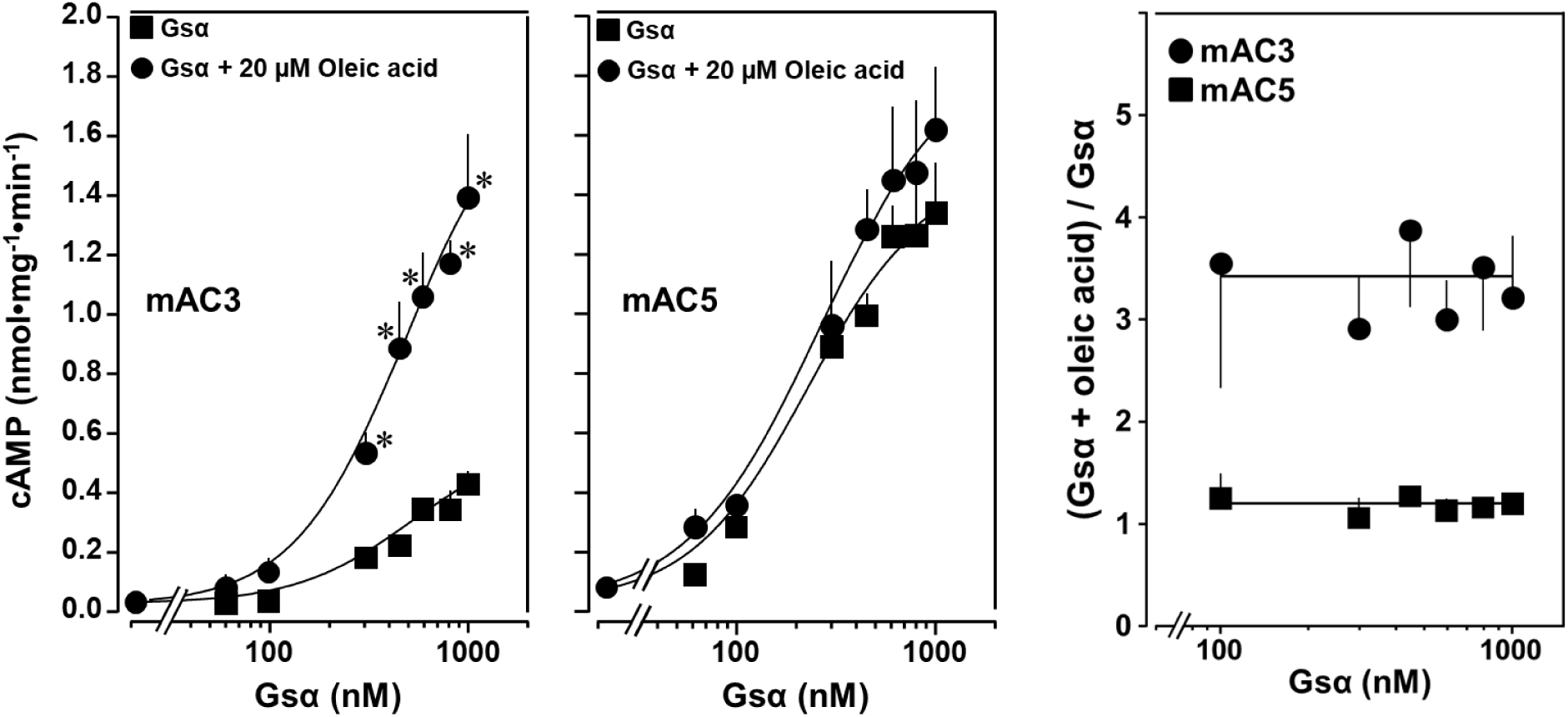
Concentration-response curves for Gsα and mAC3 and 5 activities in presence or absence of 20µM oleic acid. 20 µM oleic acid enhances Gsα stimulation of mAC3 (left) but not of mAC5 (center). **(Left)** EC_50_ of Gsα in the absence of oleic acid was 549 nM and in the presence of 20 µM oleic acid, it was 471 nM (not significant). mAC3 basal activity was 30 ± 24 pmol cAMP•mg^-1^•min^-1^. **(Center)** The EC_50_ of Gsα in the absence of oleic acid was 245 nM and in the presence of oleic acid, it was 277 nM (not significant). mAC5 basal activity was 84 ± 60 cAMP•mg^-1^•min^-1^. n=2-3, each with two technical replicates. **(Right)** (Gsα + oleic acid stimulation) / (Gsα stimulation) ratio of mAC3 and mAC5 from left and center (n = 2-3). Data are mean ± SEM, paired t test for left and center, and one-way ANOVA for right. Significances: \**P* < 0.05.

The action of oleic acid on mAC3 was linear for >25 min (appendix 2, Fig. 1). The Km of mAC3 for ATP (335 µM) was unaffected. Vmax was increased from 0.62 to 1.23 nmol cAMP/mg/min (appendix 2, Fig. 2). Oleic acid did not affect the activity of a soluble, Gsα stimulated construct formerly used for generating a C1 and C2 catalytic dimer from mAC1 and 2, ruling out spurious detergent effects (13) (appendix 2, Fig. 3). The effect of oleic acid was further evaluated by Gsα concentration response curves of mAC3 and mAC5 in presence and absence of 20 µM oleic acid (Fig. 2, left and center). For mAC3 the calculated EC_50_ of Gsα in presence and absence of oleic acid were 549 and 471 nM, respectively (not significant). Over the Gsα concentration range tested with mAC3 the enhancement of cAMP formation by 20 µM oleic was uniformly about 3.4-fold (Fig. 2 right). In the case of mAC5, Gsα stimulation was not enhanced by oleic acid (Fig. 2 center and right).

**Fig. 3.**
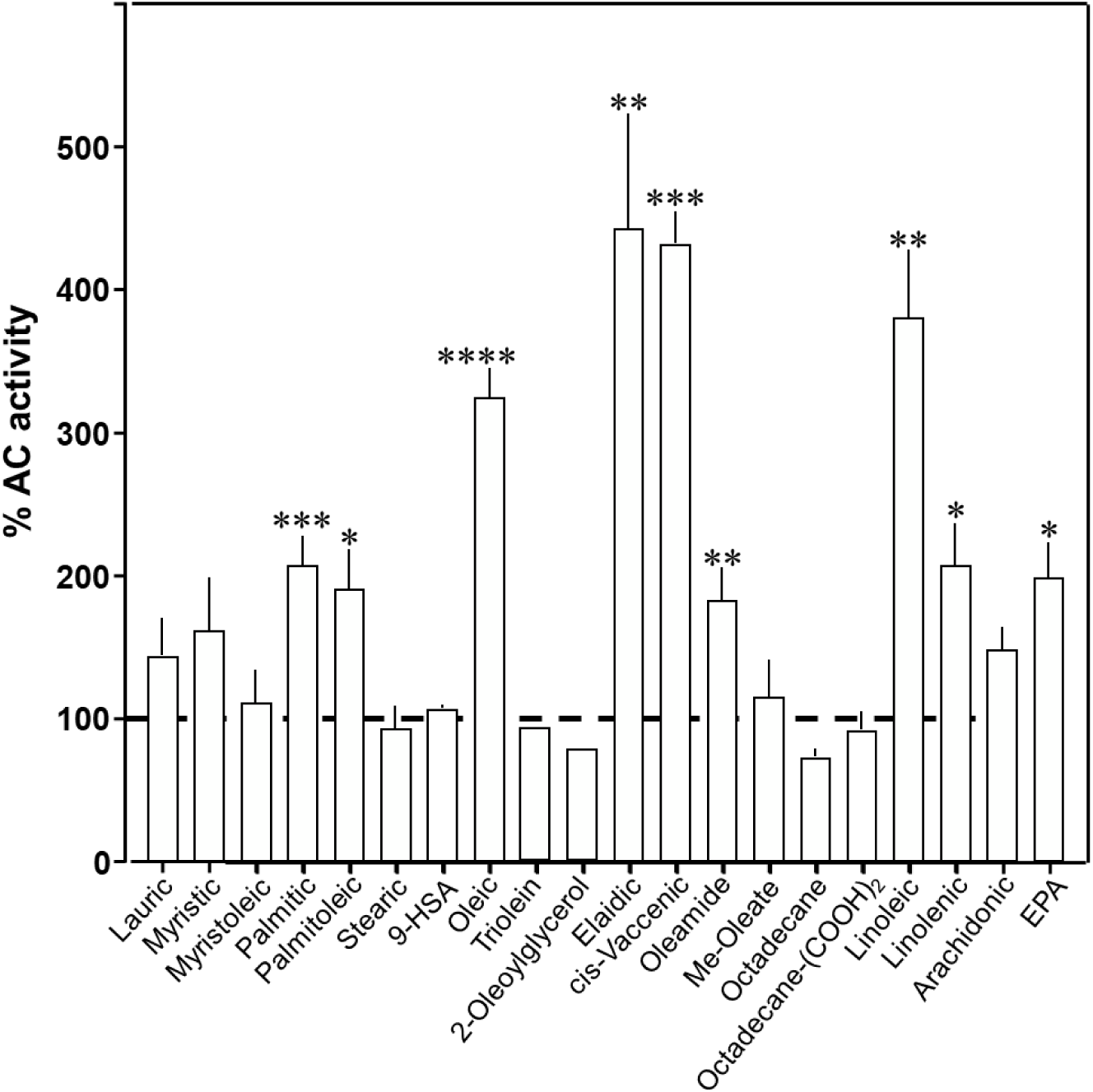
Effect of 20 µM lipids on 300 nM Gsα-stimulated mAC3. Basal and Gsα stimulated activities were 0.02 ± 0.001 and 0.17 ± 0.01 nmol cAMP•mg^-1^•min^-1^, respectively. EPA: Eicosapentaenoic acid; 9-HSA: 9-hydroxystearic acid. Data are mean ± SEM, n= 2-23. One-sample *t* tests, Significances: \**P* < 0.05, \*\**P* < 0.01, \*\*\**P* < 0.001; \*\*\*\**P* < 0.0001.

To explore the ligand space, we tested 18 aliphatic C_12_ to C_20_ lipids (Table 1; structures in appendix 2 Fig. 4). At 20 µM, elaidic, *cis*-vaccenic and linoleic acids were efficient enhancers of Gsα-stimulated mAC3 activity. Palmitic, palmitoleic, linolenic, eicosa-pentaenoic acids and oleamide were less efficacious; other compounds were inactive (Fig. 3). Notably, the saturated C_18_ stearic acid was inactive here and throughout, albeit otherwise variations in chain length, and the number, location, and conformation of double bonds were tolerated to some extent, e.g., *cis-*vaccenic, linoleic and linolenic acids. The relaxed ligand specificity was anticipated as aliphatic fatty acids are highly bendable and bind to a flexible dodecahelical protein dimer embedded in a fluid lipid membrane. The ligand space of mAC3 somewhat resembled the fuzzy and overlapping ligand specificities of the free-fatty-acid receptors 1 and 4 (25–27).

**Table 1.** List of lipids tested against mAC isoforms.

|  |
| --- |
| **Lauric (Dodecanoic) acid** |
| **Myristic (Tetradecanoic) acid** |
| **Myristoleic ((9Z)-Tetradecenoic) acid** |
| **Palmitic (Hexadecanoic) acid** |
| **Palmitoleic ((9Z)-Hexadecenoic) acid** |
| **Octadecane** |
| **1,18-Octadecanedicarboxylic acid** |
| **Stearic (Octadecanoic) acid** |
| **9-Hydroxystearic acid** |
| **Oleic ((9Z)-Octadecenoic) acid** |
| **Oleamide ((9Z)-Octadecenamide)** |
| **Methyl oleate** |
| **2-Oleoylglycerol** |
| **Triolein** |
| **Elaidic ((9E)-Octadecenoic) acid** |
| **cis-Vaccenic ((11E)-Octadecenoic) acid** |
| **Linoleic ((9Z,12Z)-Octadecadienoic) acid** |
| **Linolenic ((9Z,12Z,15Z)-Octadecatrienoic) acid** |
| **Arachidonic ((5Z,8Z,11Z,14Z)-eicosatetraenoic) acid** |
| **Eicosapentaenoic ((5Z,8Z,11Z,14Z,17Z)-eicosapentaenoic) acid** |

**Fig. 4.**
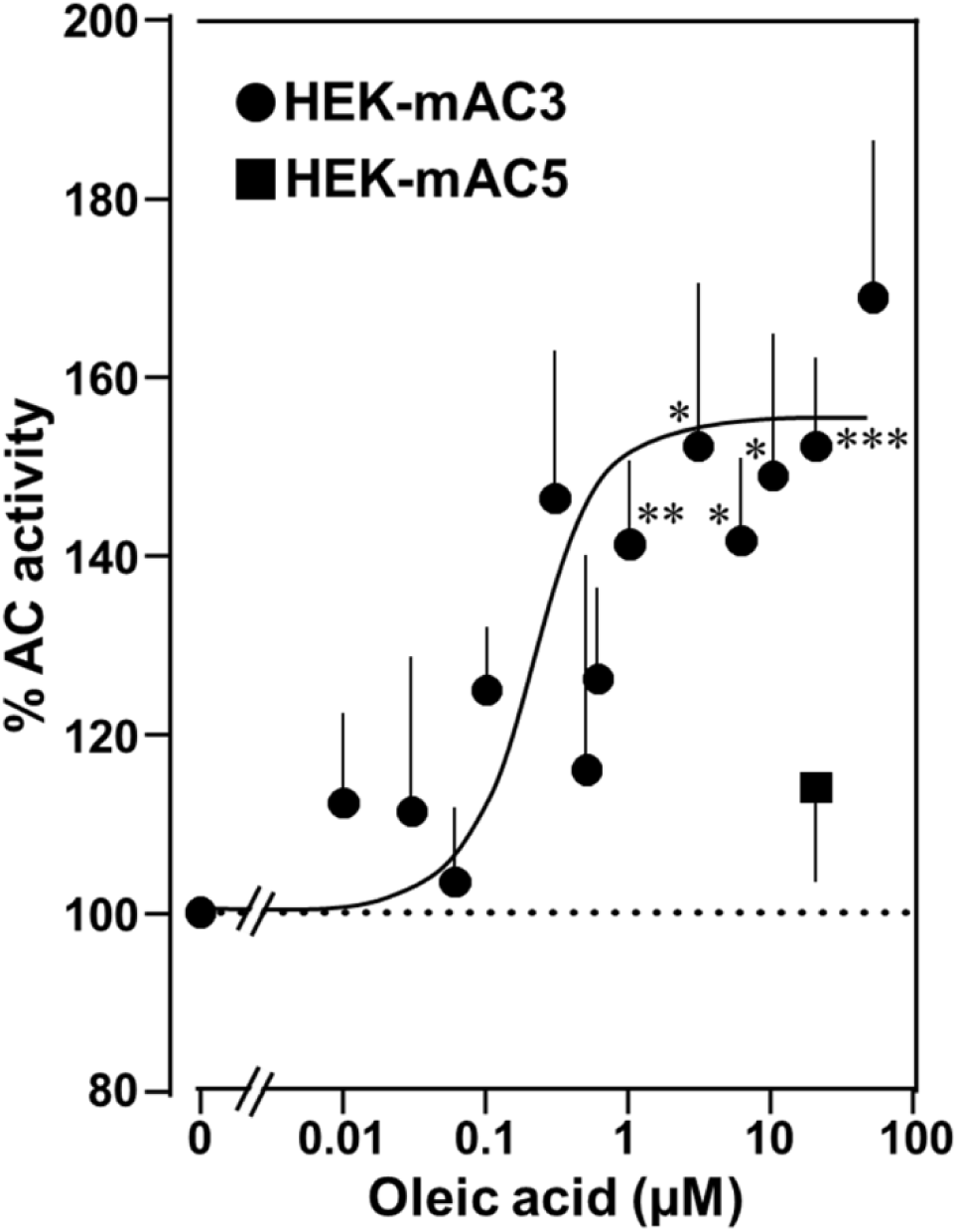
Oleic acid enhances cAMP formation in mAC3-transfected HEK293 cells. Effect of oleic acid on HEK293 cells permanently transfected with mACs 3 and 5 stimulated by 2.5 µM isoproterenol (set as 100%). Basal and isoproterenol stimulated cAMP levels of HEK-mAC3 were 0.02 ± 0.006 and 1.35 ± 0.24 and of HEK-mAC5 2.13 ± 0.69 and 2.60 ± 0.88 pmol cAMP/10000 cells, respectively. n= 3-9, carried out in technical triplicates. Data are mean ± SEM . One-sample *t* tests. Significances: \**P* < 0.05, \*\**P* < 0.01, \*\*\**P* < 0.001.

Next, the effect of oleic acid was probed *in vivo* using HEK293 cells permanently transfected with mAC3 (HEK-mAC3) or mAC5 (HEK-mAC5). Intracellular cAMP formation via Gsα was triggered via stimulation of the endogenous ß-receptor with 2.5 µM of the β-agonist isoproterenol (concentration of isoproterenol is based on a respective concentration-response curve with HEK-mAC3 cell; see appendix 2; Fig. 5). Addition of oleic acid enhanced cAMP formation in HEK-mAC3 1.5-fold (Fig. 4). Stearic acid was inactive. Under identical conditions, cAMP formation in HEK-mAC5 cells was unaffected (Fig. 4). The EC_50_ of oleic acid in HEK293-mAC3 cells was 0.5 µM, i.e., the potency appeared to be increased compared to respective membrane preparations whereas the efficiency was reduced, possibly reflecting the regulatory interplay within the cell. To exclude experimental artifacts, transfection efficiencies were tested by PCR. mAC3 and mAC5 transfections were similar (appendix 2; Fig. 6). Taken together, the results suggest that the enhancement of Gsα-stimulated mAC3 by oleic acid might be due to binding of oleic acid to or into an mAC3 membrane receptor (12, 22).

**Fig. 5.**
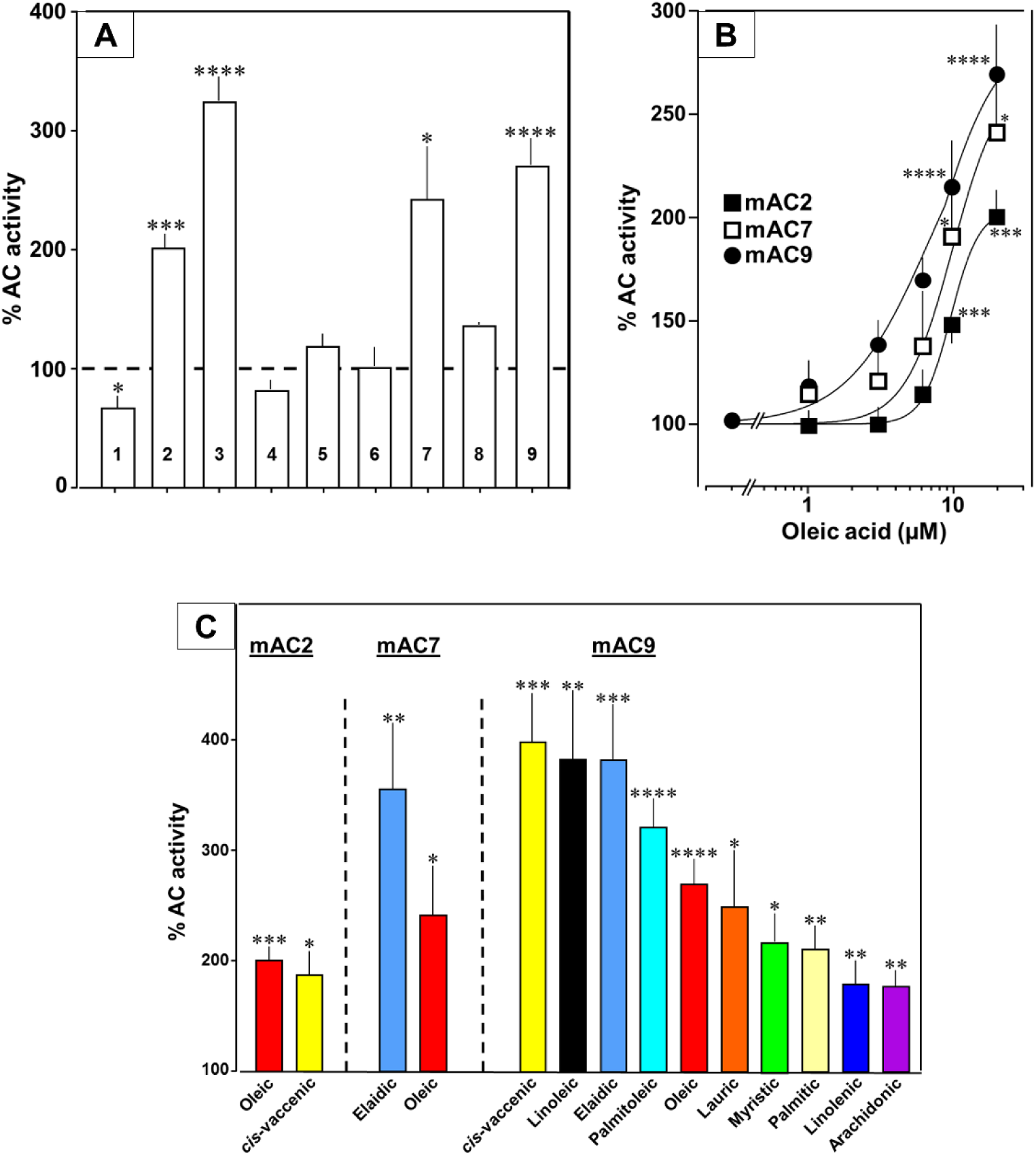
Fatty acids enhance mAC isoforms 2, 7 and 9 activities. **(A)** Effect of 20 µM oleic acid on 300 nM Gsα-stimulated mAC activities normalized to 100%. Basal and Gsα-stimulated activities for each isoform are in appendix 2 Table 1. n= 2-23. **(B)** Oleic acid activates mACs 2, 7, and 9 stimulated by 300 nM Gsα. n=7-23. **(C)** Fatty acids activating mACs 2, 7, and 9 at 20 µM. For basal and Gsα-stimulated activities, see appendix 2, Fig.’s 6-9. n= 5-15. Identical colours indicate identical compounds. Data are mean ± SEM. One-sample *t* tests were performed. Significances: \**P* < 0.05, \*\**P* < 0.01, \*\*\**P* < 0.001; \*\*\*\**P* < 0.0001.

**Fig. 6.**
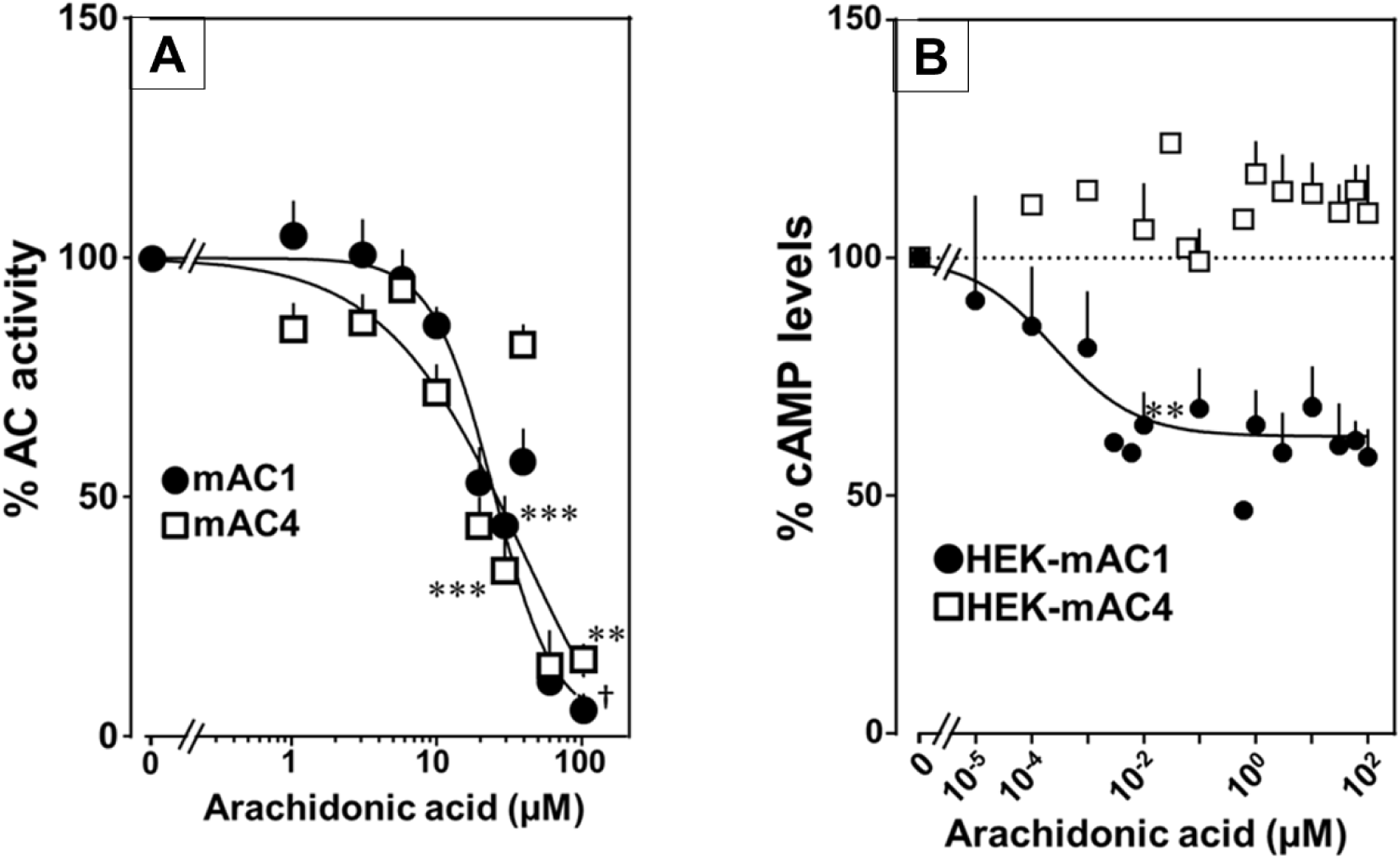
Arachidonic acid attenuates 300 nM Gsα-stimulated activities of mACs 1 and 4. **(A)** Arachidonic acid attenuates Gsα-stimulated mACs 1 and 4. Basal and Gsα stimulated activities of mAC1 were 0.12 ± 0.01 and 0.42 ± 0.03 and of mAC4 0.02 ± 0.002 and 0.14 ± 0.02 nmol cAMP•mg^-1^•min^-1^, respectively. IC_50_ of arachidonic acid for mAC1 and mAC4 were 23 and 36 μM, respectively. n= 3-9. **(B)** Effect of arachidonic acid on HEK-mAC1 and HEK-mAC4 cells. Cells were stimulated by 10 µM isoproterenol (set as 100 %) in the presence of 0.5 mM IBMX (3-isobutyl-1-methylxanthine). Basal and isoproterenol stimulated cAMP levels in HEK-mAC1 were 1.03 ± 0.15 and 1.66 ± 0.28 and in HEK-mAC4 0.20 ± 0.04 and 0.86 ± 0.24 pmol cAMP/10000 cells, respectively. IC_50_ for HEK-mAC1 was 250 pM. n= 2-11, each with three replicates. Data are mean ± SEM. One-sample *t* tests were performed. Significances: \*\**P* < 0.01, \*\*\**P* < 0.001; †*P* < 0.0001. For clarity, not all significances are indicated.

## Oleic acid enhances Gsα-stimulated mAC 2, 7, and 9 activities

Next, we examined other AC isoforms with oleic acid as a ligand. 20 µM oleic acid significantly enhanced Gsα-stimulated activities of isoforms 2, 7, and 9, mAC1 was slightly attenuated, and isoforms 4, 5, 6, and 8 were unaffected (Fig. 5A).

Concentration-response curves were carried out for mACs 2, 7, and 9 (Fig. 5B). The EC_50_ of oleic acid were 8.6, 6.7 and 7.8 µM, respectively, comparable to that determined for mAC3. Exploration of the ligand space for mACs 2, 7 and 9 with the panel of 18 aliphatic lipids uncovered more active lipids (Fig. 5C). In the case of mAC2, 20 µM *cis*-vaccenic acid doubled cAMP formation (EC_50_ 10.6 µM) while other compounds were inactive (Fig. 5C and for additional concentration-response curves see appendix 2 Fig. 7). For mAC7 the EC_50_ of elaidic was 9.7 µM (concentration-response curve see appendix 2 Fig. 8). The range of potential ligands for mAC9 was more comprehensive: 3-4-fold enhancement was observed with 20 µM palmitoleic, oleic, elaidic, *cis*-vaccenic, and linoleic acid. With 20 µM myristic, palmitic, palmitoleic, linolenic, and arachidonic acid 1.5-2 fold enhancements were observed (Fig. 5C and for concentration response curves see appendix 2 Fig.s 9 and 10).

**Fig. 7.**
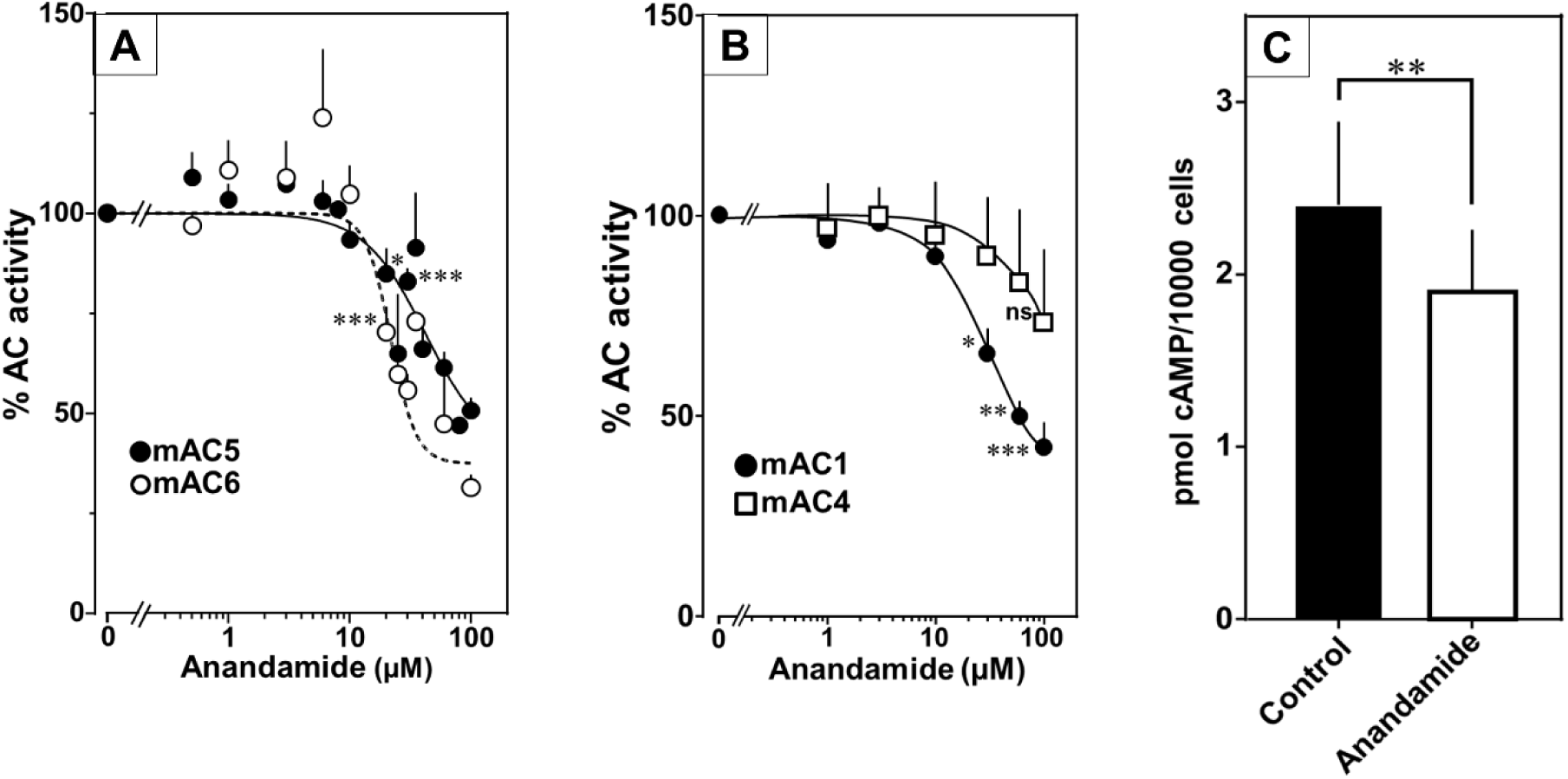
Anandamide attenuates 300 nM Gsα-stimulated activities of mACs 1, 4, 5 and 6. **(A)** Effect of anandamide on Gsα-stimulated mAC5 and 6. Basal and Gsα activities of mAC5 were 0.05 ± 0.01 and 0.98 ± 0.12 and of mAC6 0.05 ± 0.01 and 0.78 ± 0.12 nmol cAMP•mg^-1^•min^-1^, respectively. IC_50_ of anandamide were 42 and 22 μM, respectively. n= 3-32. **(B)** Anandamide attenuates mAC1 but not mAC4 stimulated by Gsα. Basal and Gsα stimulated activities of mAC1 were 0.12 ± 0.01 and 0.40 ± 0.03 and of mAC4 0.02 ± 0.002 and 0.15 ± 0.02 nmol cAMP•mg^-1^•min^-1^, respectively. IC_50_ for mAC1 was 29 μM. n= 3-4, each with two technical replicates. **(C)** Effect of anandamide on 2.5 µM isoproterenol stimulated HEK-mAC5. Basal and isoproterenol stimulated cAMP levels of HEK-mAC5 were 1.8 ± 0.22 and 2.4 ± 0.48 pmol cAMP/10000 cells, respectively. The control bar represents 2.5 µM isoproterenol stimulation alone. n=5-6, each with three technical replicates. IC_50_ of anandamide was 133 µM. Data are mean ± SEM. One-sample *t* tests (A-B) and paired *t* test (C) were performed. Significances: ns: not significant *P* > 0.05;\**P* < 0.05, \*\**P* < 0.01, \*\*\**P* < 0.001. For clarity, not all significances are indicated.

**Fig. 8.**
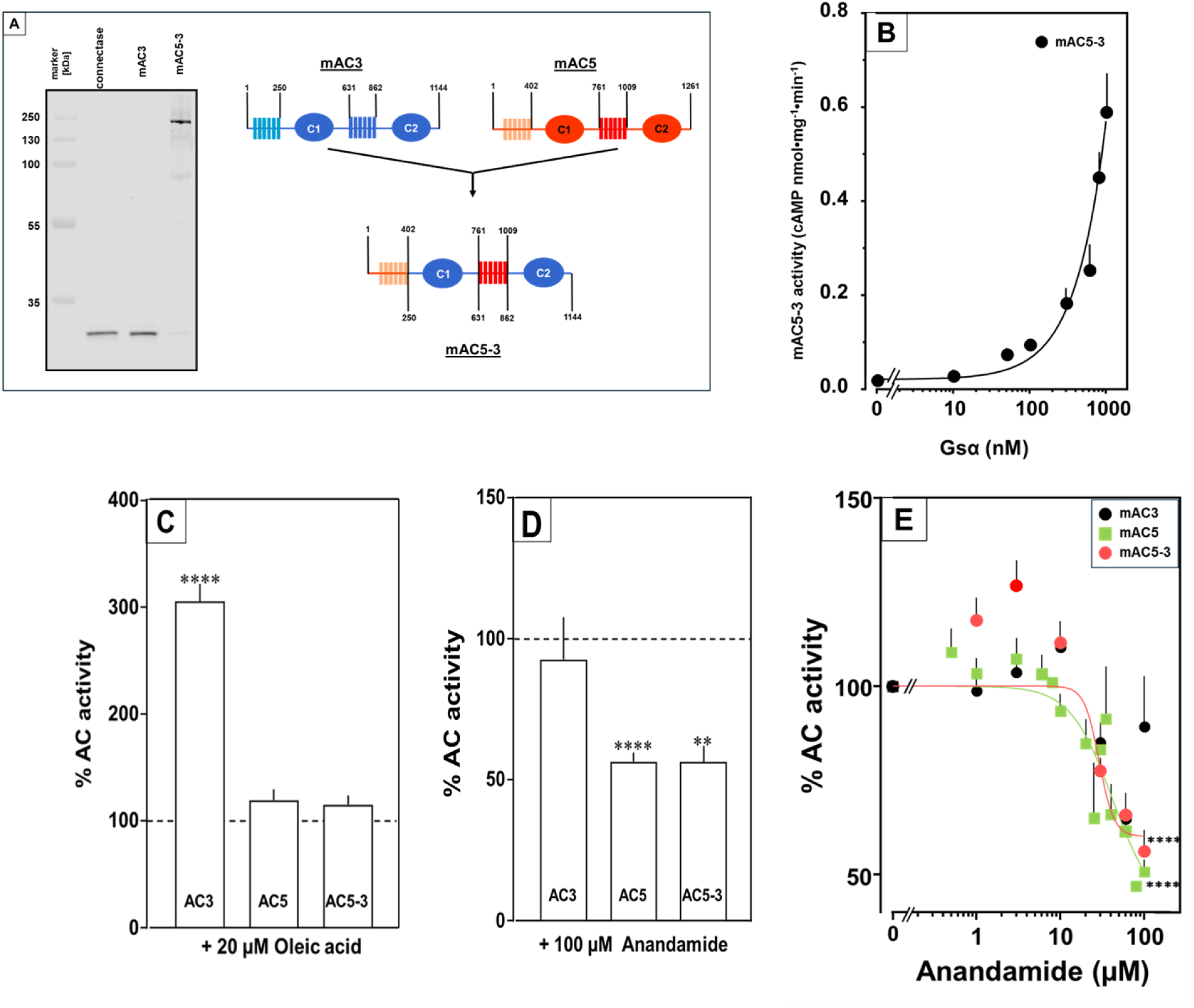
Receptor properties are exchangeable between mAC isoforms. **(A, left)** Detection of AC5_(membr)_-AC3_(cat)_ receptor chimeras. AC5_(membr)_-AC3_(cat)_ (AC5-3) (36) was expressed in HEK293 cells with an N-terminal tag for labeling with the protein ligase Connectase. The membrane preparation was incubated with fluorophore-conjugated Connectase and separated by SDS-PAGE. A fluorescence scan of the gel detects AC5_(membr)_-AC3_(cat)_ (right), the reagent (fluorophore-conjugated Connectase) is detected when using HEK293 membrane (middle) or a buffer control (left); (**A, right)** Design of the chimeric AC5-3 construct. Numbers are amino acid positions in mAC3 and 5, respectively. **(B)** Gsα concentration response curve of mAC5-3. Basal activity for mAC5-3 was 0.02 pmol cAMP•mg^-1^•min^-1^. Error bars denote SEM of n=3, each with two technical replicates. **(C)** Effect of 20 µM oleic acid on 300 nM Gsα-stimulated mACs 3, 5, and 5-3. Basal and Gsα activities of mACs 3, 5, and 5-3 were 0.02 ± 0.003 and 0.11 ± 0.02, 0.05 ± 0.01 and 0.98 ± 0.12, and 0.01 ± 0.004 and 0.2 ± 0.02 nmol cAMP•mg^-1^•min^-1^, respectively. n=7-33. **(D)** Effect of 100 µM Anandamide on 300 nM Gsα-stimulated mACs 3, 5, and 5-3. Basal and Gsα activities of mACs 3, 5, and 5-3 were 0.02 ± 0.002 and 0.19 ± 0.02, 0.05 ± 0.01 and 0.98 ± 0.12 and 0.02 ± 0.003 and 0.23 ± 0.04 nmol cAMP•mg^-1^•min^-1^, respectively. n=6-9. IC_50_ for mAC5 and mAC5-3 were 42 and 29 µM, respectively. **(E).** Exchange of TM domains transfers anandamide effect on mAC3. Basal and Gsα stimulated activities of mAC3 were 0.02 ± 0.002 and 0.12 ± 0.02 nmol cAMP•mg^-1^•min^-1^, respectively. Basal and Gsα stimulated activities of mAC5 were 0.05 ± 0.005 and 0.98 ± 0.12 nmol cAMP•mg^-1^•min^-1^, respectively. Basal and Gsα stimulated activities of mAC5-3 were 0.02 ± 0.002 and 0.22 ± 0.03 nmol cAMP•mg^-1^•min^-1^, respectively. Calculated IC_50_ concentrations of Anandamide for mAC5 and mAC5-3 were 42 and 29 µM, respectively. Data are mean ± SEM. One-sample *t* tests. Significances: \*\**P* < 0.01; \*\*\*\**P* < 0.0001.

**Fig. 9.**
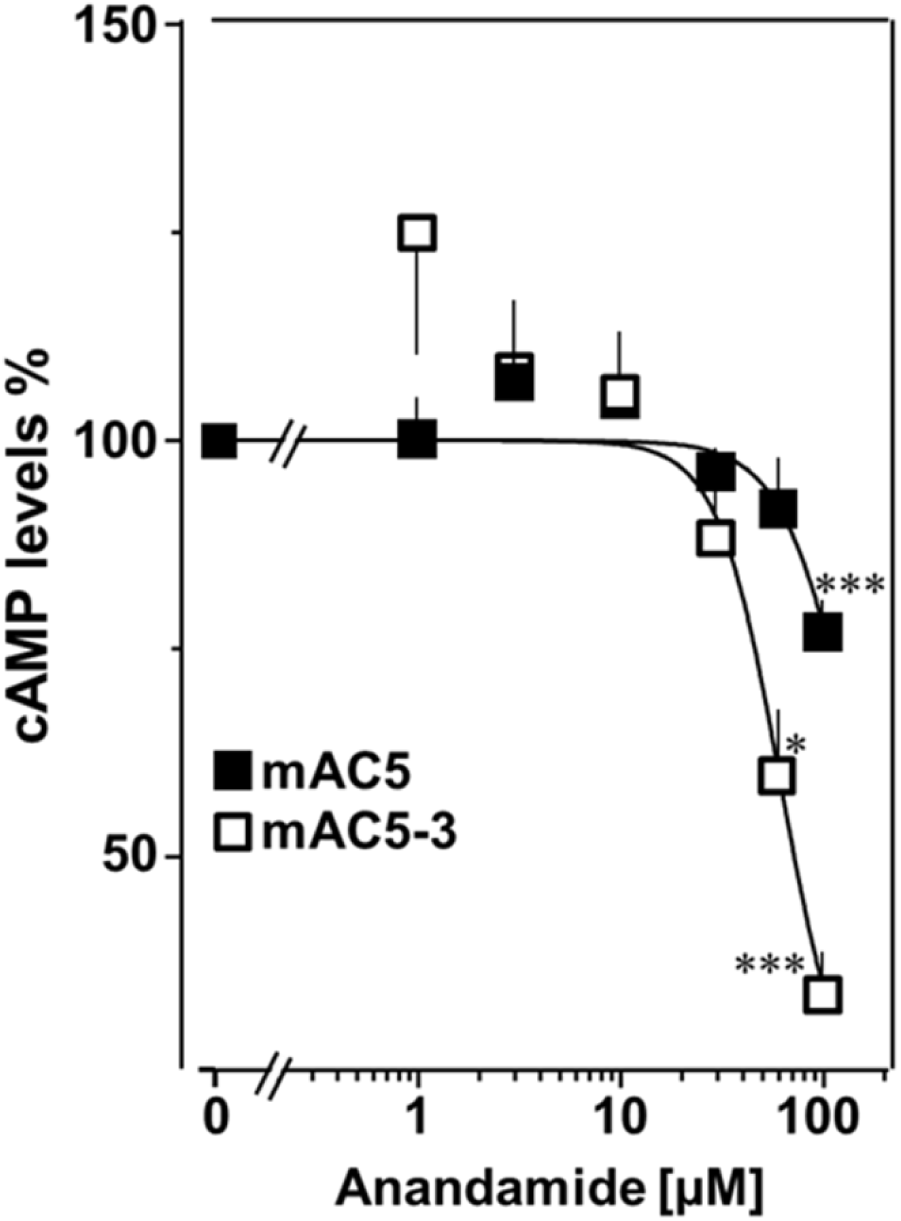
mAC5 – mAC3 receptor-transfer analyzed in HEK293 cells. Effect of anandamide on HEK-mAC5 and HEK-mAC5-3 cells stimulated by 2.5 µM isoproterenol (set as 100 %). Basal and isoproterenol stimulated cAMP levels in HEK-mAC5 were 1.80 ± 0.22 and 2.29 ± 0.39 and in HEK-mAC5-3 (+ 0.5 mM IBMX) 0.17 ± 0.02 and 3.11 ± 0.55 pmol cAMP/10000 cells, respectively. n= 4-11. IC_50_ for HEK-mAC5 and HEK-mAC5-3 were 133 and 60 μM, respectively. Anandamide had no effect on basal activity of HEK-mAC5 and stimulated HEK-mAC3 cells in concentrations up to 100 µM (data not shown). Data are mean ± SEM. One-sample *t* tests were performed. Significances: \**P* < 0.05; \*\*\**P* < 0.001.

**Fig. 10.**
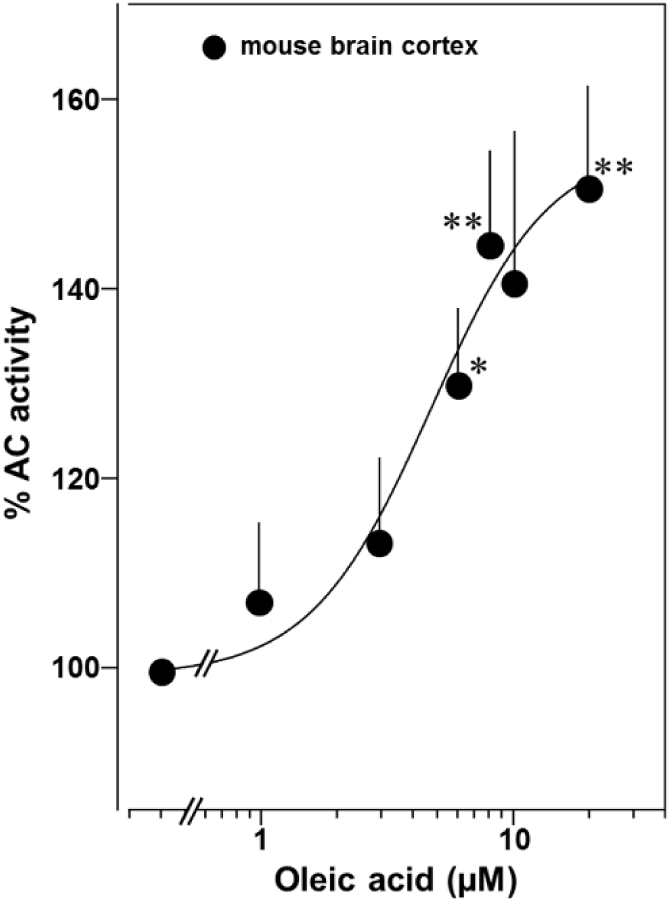
Oleic acid concentration-dependently potentiates mAC activity in brain cortical membranes from mouse. Basal and 300 nM Gsα activities were 0.4 ± 0.1 and 2.7 ± 0.7 nmol cAMP•mg^-1^•min^-1^, respectively. N= 4-6. One-sample *t* test: \**P* < 0.05; \*\**P* < 0.01 compared to 100% (300 nM Gsα stimulation).

## Arachidonic acid and anandamide inhibit Gsα-stimulated activities of mAC1, 4, 5 and 6

Testing the panel of lipids at 20 µM with mAC isoforms 1, 4, 5, 6, and 8 we found that isoforms 1 and 4 were significantly attenuated by arachidonic acid, and somewhat less by palmitoleic acid. Other lipids had no effect (for bar plots and dose response curves see appendix 2 Fig. 11-13). Of note, eicosa-pentaenoic acid which resembles arachidonic acid but for an additional *cis*-Δ^-17^ double bond had no effect on mAC activities (appendix 2 Fig. 11, 12). Concentration-response curves for arachidonic acid with 300 nM Gsα-stimulated mAC1 and 4 yielded IC_50_ of 23 and 36 µM, respectively, i.e., about two-fold higher compared to the EC_50_ of enhancing ligands (Fig. 6A).

Next, we examined whether arachidonic acid attenuates mAC1 and 4 in intact HEK 293 cells. Surprisingly, cAMP formation in HEK-mAC1 cells stimulated by 10 µM isoproterenol was attenuated by arachidonic acid with high potency (IC_50_ = 250 pM), i.e., with higher potency compared to data with membranes prepared from the same cell line. In contrast, mAC4 activity examined under identical conditions was not attenuated (Fig. 6B). Currently, we are unable to rationalize these discrepancies. Possibly, mAC4 has another, more specific lipid ligand which is needed in *in vivo*. In general, the enhancing and attenuating effects bolster the hypothesis of specific receptor-ligand interactions and divergent intrinsic activities for different ligands. Of note, it was reported that arachidonic acid at concentrations up to 1 mM inhibits AC activity in brain membrane fractions and that essential fatty acid deficiency effects AC activity in rat heart (28, 29).

At this point we were lacking ligands for mACs 5, 6, and 8 (appendix 2 Fig. 14-16). Possibly, the negative charge of the fatty acid headgroups might impair receptor interactions. A neutral lipid neurotransmitter closely related to arachidonic acid is arachidonoylethanolamide (anandamide) (30). Indeed, anandamide attenuated 300 nM Gsα-stimulation of mAC5 and 6 with IC_50_ of 42 and 23 µM, respectively, i.e., comparable to the effect of arachidonic acid on mACs 1 and 4, and distinctly less potently than the ligands for mAC 2, 3, 7, and 9 (Fig. 7A).

mACs 5 and 6 may thus represent new targets for anandamide which is part of a widespread neuromodulatory system (31). The concentrations of arachidonic acid and anandamide required may be achieved *in vivo* by local biosynthesis and degradation. An interfacial membrane-embedded phosphodiesterase cleaves the phosphodiester bond of the membrane lipid *N-*arachidonoyl-ethanolamine-glycerophosphate releasing anandamide into the extracellular space (30, 32, 33). The lipophilicity and lack of charge should enable it to diffuse readily. Whether the mACs and this biosynthetic phosphodiesterase colocalize or associate with its target mACs is unknown. Degradation of anandamide is by a membrane-bound amidase, generating arachidonic acid and ethanolamine (34). Therefore, we examined whether anandamide at higher concentrations might also affect mAC1 and 4. In fact, anandamide significantly attenuated Gsα-stimulated mAC1, but distinctly not mAC4 (Fig. 7B). The IC_50_ of anandamide for mAC1 was 29 µM.

We also tested whether anandamide attenuated cAMP formation *in vivo* using HEK-mAC5 cells primed by 2.5 µM isoproterenol (Fig. 7C). 100 µM Anandamide attenuated cAMP formation by only 23 % in HEK293-mAC5 cells, the effect was significant (*P* < 0.01). At this point, we were unable to identify a ligand for mAC8, presumably another lipid.

## Receptor properties are exchangeable between mAC isoforms 3 and 5

To unequivocally validate specific mAC-ligand-receptor interactions and regulation we generated a chimera in which the enhancing membrane domains of mAC3, i.e., mAC3-TM1 and TM2, were substituted by those of mAC5 (design in Fig. 8A, right). The intention was to obtain a chimera, mAC5_(membr)_-AC3_(cat),_ with a loss of receptor function, i.e., no enhancement by oleic acid, and a gain of another receptor function, i.e., attenuation of activity by anandamide. Successful expression and membrane insertion of the chimera in HEK293 cells was demonstrated by specific conjugation to Cy5.5 fluorophore, using the protein ligase connectase (Fig. 8A, left (35)). cAMP synthesis of isolated membranes from these cells was stimulated up to 10-fold by addition of 300 nM Gsα, comparable to membranes with recombinant mAC3 or mAC5 proteins (Fig. 8B). mAC activity in the mAC5_(membr)_-AC3_(cat)_ chimera was not enhanced by oleic acid, i.e., loss of receptor function, but was attenuated by anandamide, i.e., gain of receptor function (Fig. 8C and D). The attenuation was comparable to results obtained with mAC5 membranes, i.e., IC_50_ was 29 µM mAC5_(membr)_-AC3_(cat)_ (Fig. 8E) compared to 42 µM for mAC5. This means that the attenuating receptor property of mAC5 was successfully grafted onto the mAC3-catalytic dimer. We take this to support the hypothesis that the mammalian mAC membrane domains operate as receptors using lipid ligands. The data virtually rule out unspecific lipid effects such as disturbance of membrane integrity by intercalation and surfactant or detergent effects. In addition, the data demonstrated that the signal most likely originates from the receptor entity and is transmitted through the subsequent linker regions to the catalytic dimer.

The findings were further substantiated *in vivo* using HEK293-mAC5_(membr)_-AC3_(cat)_ cells. cAMP formation primed by 2.5 µM isoproterenol was attenuated by anandamide in HEK293-mAC5_(membr)_-AC3_(cat)_ cells by 66%, (Fig. 9), i.e., a gain of function which remarkably exceeded the anandamide attenuation in HEK293-mAC5 cells of 23 %. In HEK293-mAC5_(membr)_-AC3_(cat)_ cells oleic acid was ineffective, i.e., loss of function (data not shown). The results bolster the notion that mAC isoforms are receptors with lipids as ligands.

Lastly, we prepared membranes from mouse brain cortex in which predominantly mAC isoforms 2, 3 and 9 are expressed, isoforms with demonstrated enhancement of Gsα stimulation by oleic acid (37). In cortical membranes 20 µM oleic acid enhanced Gsα stimulated cAMP formation 1.5-fold with an EC_50_ of 5 μM, almost identical to the one determined for mAC2, 3, 7 and 9 (Fig. 10). This suggests that mACs in brain cortical membranes are similarly affected by fatty acids.

## Discussion

In the past, the biology of the two membrane anchors of mACs, highly conserved in an isoform-specific manner, remained unresolved. The theoretical possibility of a receptor function of these large hexahelical anchor domains, each comprising 150-170 amino acids, was considered unlikely (14). Our data are a transformative step toward resolving this issue and introduce lipids as critical participants in regulating cAMP biosynthesis in mammals. The first salient discovery is the identification of the membrane domains of mACs as a new class of receptors for chemically defined ligands which set the level of stimulation by the GPCR/Gsα system. This conclusion is based on *(i)* the dodecahelical membrane domains of the nine mAC receptors have distinct, conserved isoform-specific sequences for the TM1 and TM2 domains (12); *(ii)* the receptors have distinct ligand specificities and affinities in the lower micromolar range; *(iii)* isoform dependently ligands either enhance or attenuate Gsα-stimulated mAC activities; *(iv)* receptor properties are transferable between isoforms by interchanging membrane domains; *(v)* isoproterenol-stimulated formation of cAMP *in vivo* is affected by addition of extracellular ligands. *(vi)* Gsα-stimulated cAMP formation in mouse cortical membranes is enhanced by oleic acid. Therefore, the results establish a new class of receptors, the membrane domains of mACs, with lipids as ligands. The data question the utility of the currently used mAC sub-classification, which groups mAC1, 3, 8, mAC2, 4, 7, mACs 5, 6, and mAC9 together (2). At this point, mAC 1, 4, 5, and 6, which are ligand-attenuated, may be grouped together and a second group may consist of isoforms 2, 3, 7, and 9 which are ligand-enhanced. Our data do not contradict earlier findings concerning regulation of mACs via GPCRs, cellular localization of mAC isoforms or regional cAMP signaling (2). Instead, the data reveal a completely new level of regulation of cAMP biosynthesis in which two independent modalities of signaling, i.e., direct, tonic lipid signaling and indirect phasic signaling via the GPCR/Gsα circuits intersect at the crucial biosynthetic step mediated by the nine mAC isoforms.

The second important finding is the observation that the extent of enhancement of mAC3 activity by 20 µM oleic acid is uniform up to 1000 nM Gsα (Fig. 2 right). We suppose that in mAC3 the equilibrium of two differing ground states favors a Gsα unresponsive state and the effector oleic acid concentration-dependently shifts this equilibrium to a Gsα responsive state (19). In contrast, the equilibrium of ground states of mAC5 probably is opposite, i.e. the one accessible to Gsα stimulation predominates and stimulation by Gsα is maximal. In addition, oleic acid has little effect because the mAC5 receptor domain does not bind oleic acid (Fig. 1D, right, and Fig. 2, center). A ligand for mAC5, e.g., anandamide or arachidonic acid, likely shifts the equilibrium of ground states to a Gsα unresponsive state and inhibits stimulation. The biological balance of ground states appears to be an intrinsic property which is isoform-specifically imprinted in mACs. Probably, it defines a major element of regulation and enables distinct inhibitory or stimulatory inputs by extracellular lipid ligands. The ground states probably are separated by a low transition energy and are stabilized by receptor occupancy. Hitherto available structures required Gsα and/or forskolin for stabilization and probably did not capture different ground states (14, 16, 17, 38–40). Mechanistically, tonic levels of lipid ligands affect the ground states and thus set the bounds of cAMP formation elicited by phasic GPCR/Gsα-stimulation. As such lipid signaling through the mAC membrane receptors appears to represent a higher level of a systemic regulatory input based on constant monitoring the physiological and nutritional status of an organism.

Lipid signaling is much less characterized than solute signaling (41). Most of the highly functionalized ligands for GPCRs are storable in vesicles and the release, inactivation and removal are strictly controlled. On the other hand, the very nature of lipids, i.e., high flexibility of aliphatic chains, low water solubility, propensity for nonspecific protein binding, membrane permeability and potential effects on membrane fluidity complicate discrimination between extra- and intracellular lipid actions (26). Yet, viewed from an evolutionary perspective, lipids possibly are ideal primordial signaling molecules because for the emergence of the first cell lipids were required to separate an intra- and extracellular space. Conceivably, lipids derived from membrane lipids were used for regulatory purposes early-on. In association with the evolution of bacterial mAC progenitors lipid ligands may have persisted in evolution and regulation by GPCR/Gsα in metazoans was acquired and expanded later.

The concentrations of free fatty acids in serum or interstitial fluid usually are rather low, mostly below the EC_50_ concentrations determined in this study (27, 42, 43). This raises the question of the origin of lipid ligands under physiological conditions. It is well known that cell membranes are highly dynamic (29, 44). Within limits, cell morphology and lipid composition are in constant flux to accommodate diverse functional requirements.

Remodeling of cell membranes is accomplished by targeted phospholipid biosynthesis and by regulated lipolysis of membrane lipids, e.g., by phospholipase A_2_, mono- and diacylglycerol lipases or lipoprotein lipases (45). Therefore, a speculative possibility is that lipid ligands are acutely and locally generated directly from membrane lipids (33, 46). Such regulation probably happens at the level of individual cells, cellular networks, complex tissues and whole organs. Additional potential lipid sources available for ligand generation may be, among others, lipids in nutrients (29), exosomes present in blood, serum lipids, chylomicrons, blood triglycerides and lipids originating from the microbiome. On the one hand, the discovery of lipids as ligands for the mAC receptors broadens the basis of regulation of cAMP biosynthesis with potentially wide-ranging consequences in health and disease. On the other hand, the data pose the challenge to identify how the tonic signal is generated and regulated

## MATERIALS AND METHODS

### Reagents and materials

ATP, creatine kinase, and creatine phosphate were from Merck. Except for lauric acid (Henkel) and 1,18-Octadecanedicarboxylic acid (Thermofisher Scientific), lipids were from Merck. 10 mM stock solutions were prepared in analytical grade DMSO and kept under nitrogen. For assays the stock solution was appropriately diluted in 20 mM Mops buffer, pH 7.5, suitable to be added to the assays resulting in the desired final concentrations. The final DMSO concentrations in *in vitro* and *in vivo* assays were maximally 1%, a concentration without any biochemical effect as checked in respective control incubations. The constitutively active GsαQ227L mutant protein was expressed and purified as described earlier (47–49).

### General Experimental Procedures

For HPLC analysis, a Waters HPLC system (1525 pump, 2996 photodiode array detector, 7725i injector, 200 series PerkinElmer vacuum degasser) was used. Solvents were HPLC or LC-MS grade from Merck-Sigma. One-dimensional ^1^H and ^13^C NMR spectra were recorded on a 400 MHz Bruker AVANCE III NMR spectrometer equipped with a 5 mm broadband SmartProbe and AVANCE III HD Nanobay console. Spectra were recorded in methanol-*d*_4_ and calibrated to the residual solvent signal (*δ*_H_ 3.31 and *δ*_C_ 49.15 ppm).

### Lung tissue extraction and fractionation

1.24 kg bovine lung was minced in a meat grinder, then mixed and homogenized with 1.2 L 50mM MOPS, pH 7.5, in a Waring blender (4°C) resulting in 2.3 L homogenate. It was centrifuged (30 min, 4°C, 7200×g) resulting in 1.2 L supernatant. The pH was adjusted to 1 using 7% HCl. Equal volumes of CH2Cl2/MeOH (2:1) were mixed with the supernatant in a separatory funnel and shaken vigorously. Centrifugation was at 5300×*g* for 30 min. The lower organic CH2Cl2 layer was recovered and the solvent was evaporated yielding 2 g of dried material. It was dissolved in 100 ml petroleum ether and subjected to normal-phase silica gel vacuum liquid chromatography (60 H Supelco). The column was eluted stepwise with solvents of increasing polarity from 90:10 petroleum ether/EtOAc to 100% EtOAc, followed by 100% MeOH. 17 fractions (A-Q) of 300 mL were collected and dried. Fraction E (eluting at 40:60 petroleum ether/EtOAc) was analysed by RP-HPLC using a linear MeOH/H_2_O gradient from 80:20 to 100:0 (0.1% TFA) for 15 min, followed by 100:0 for 30 min (Knauer Eurosphere II C18P 100-5, 250 x 8 mm, 1.2 mL/min flow rate, UV-absorbance monitored at 210 nm) to yield five subfractions; E1-E5. Fraction E2 was analysed by ^1^H- and ^13^C-NMR which indicated the presence of aliphatic lipids and fatty acids (Fig. S2).

### GC-MS analysis

Fraction E2 was analysed by GC-MS. Acids were acid trimethylsilylated using *N,O*-bis(trimethylsilyl)trifluoroacetamide + trimethylchlorosilane (BSTFA + TMCS, 99:1 v/v). The mixture was heated for 2 h at 90°C. After cooling and clearing the sample was transferred into a GC vial in 200 µL hexane.

An Agilent Technologies GC system (8890 gas chromatograph and 5977B mass spectrometer equipped with a DB-HP5MS UI column, 30 m x 0.25 mm, film thickness of 0.25 µm) was used. Injection volume was 1 µL. The temperature was kept at 100 C for 5 min, and then increased at 53°C/min to 240 C. The rate was decreased to 3 C/min to reach 305 C. Carrier gas was He_2_ (99.9%; 1.2 mL/min). Ionization was with 70 eV and MS spectra were recorded for a mass range *m/z* 35-800 for 35 min. Compounds were identified by comparing the spectra with those in the NIST library. Individual compound content is given as a relative % of the total peak area.

### Plasmid construction and protein expression

hAC sequences were from NCBI were: ADCY1: NM_021116.3; ADCY2: NM_020546.2; ADCY3: NM_004036.4; ADCY4: NM_001198568.2; ADCY5: NM_183357.2; ADCY6: NM_015270.4; ADCY7: NM_001114.4; ADCY8: NM_001115.2; ADCY9: NM_001116.3.

Human mAC genes were from GenScript and fitted with a C-terminal FLAG-tag. The chimera mAC5(TM)_mAC3(cat) had an N-terminal connectase-tag, MPGAFDADPLVVEIAAAGA, followed by AC5(1–402)_AC3(250–631)_AC5(761–1009)_AC3(862–1144). The gene was synthesized by GenScript. HEK293 cells were maintained in DMEM with 10% FBS at 37°C and 5% CO_2_. Transfection with AC plasmids was with PolyJet (SignaGen, Frederick, MD, USA). Permanent cell lines were generated by selection for 7 days with 600 µg/mL G418 and maintained with 300 µg/mL For membrane preparation, cells were tyrpsinized, collected by centrifugation (3000xg, 5 min) and lysed and homogenized in 20 mM HEPES, pH 7.5, 1 mM EDTA, 2 mM MgCl_2_, 1 mM DTT, one tablet of cOmplete, EDTA-free (per 50 mL) and 250 mM sucrose by 20 strokes in a potter homogenizer on ice. Debris was removed (5 min at 1000xg, 0°C), membranes were collected at 100,000xg, 60 min at 0°C, suspended and stored at -80°C in 20 mM MOPS, pH 7.5, 0.5 mM EDTA, 2 mM MgCl_2_. Membrane preparation from mouse brain cortex was according to (19, 50). Three cerebral cortices were dissected and homogenized in 4.5 ml cold 48 mM Tris-HCl, pH 7.4, 12 mM MgC1_2_, and 0.1 mM EGTA with a Polytron hand disperser (Kinematica AG, Switzerland). The homogenate was centrifuged for 15 min at 12000xg at 4°C. The pellet was washed once with 5 mL 1 mM KHCO_3_. The final suspension in 2 mL 1 mM KHCO_3_ was stored in aliquots at -80 C.

### DNA extraction

DNA from 1×10^6^ cells of permanently transfected and non-transfected HEK293 cells was extracted using the High Pure PCR Template Preparation Kit (Roche) according to the manufacturer’s instructions. DNA concentrations were determined at 260 nm using a sub-microliter cell (IMPLEN) in a P330 NanoPhotometer (IMPLEN). Elution buffer (Roche) was used for blanks.

### Polymerase chain reaction

100 ng of template DNA was mixed with 0.5 µM Forward primer and 0.5 µM Reverse primer. 12.5 µL 2X KAPA2G Fast (HotStart) Genotyping Mix with dye and water was added, total reaction volume 25 µL according to the KAPA2G Fast HotStart Genotyping Mix kit (Roche) protocol. PCR followed the cycling protocol in a Biometra T3000 thermocycler:

| Step | Temperature | Duration | Cycles |
| --- | --- | --- | --- |
| Initial denaturation | 95 C | 3 min | 1 |
| Denaturation | 95 C | 15 sec | 35 |
| Annealing | 60 C | 15 sec |  |
| Extension | 72 C | 30-60 sec* |  |
| Final extension | 72 C | 2-4 min* | 1 |

\*Extension and final extension times were adjusted to the expected amplicon length.

The PCR products were directly loaded on a 1.5% agarose gel. A 1 kb DNA ladder (New England Bio Labs #N3232S) was mixed with Gel Loading Dye Purple 6X (New England Biolabs #B7024S) and water. After running the gel for 15-20 minutes at 90 V in 1X TAE buffer, the gel was stained in an Ethidium bromide bath and left running for another 10-20 minutes. The gels were then evaluated under UV light in a UVP GelStudio PLUS (Analytik Jena) gel imager.

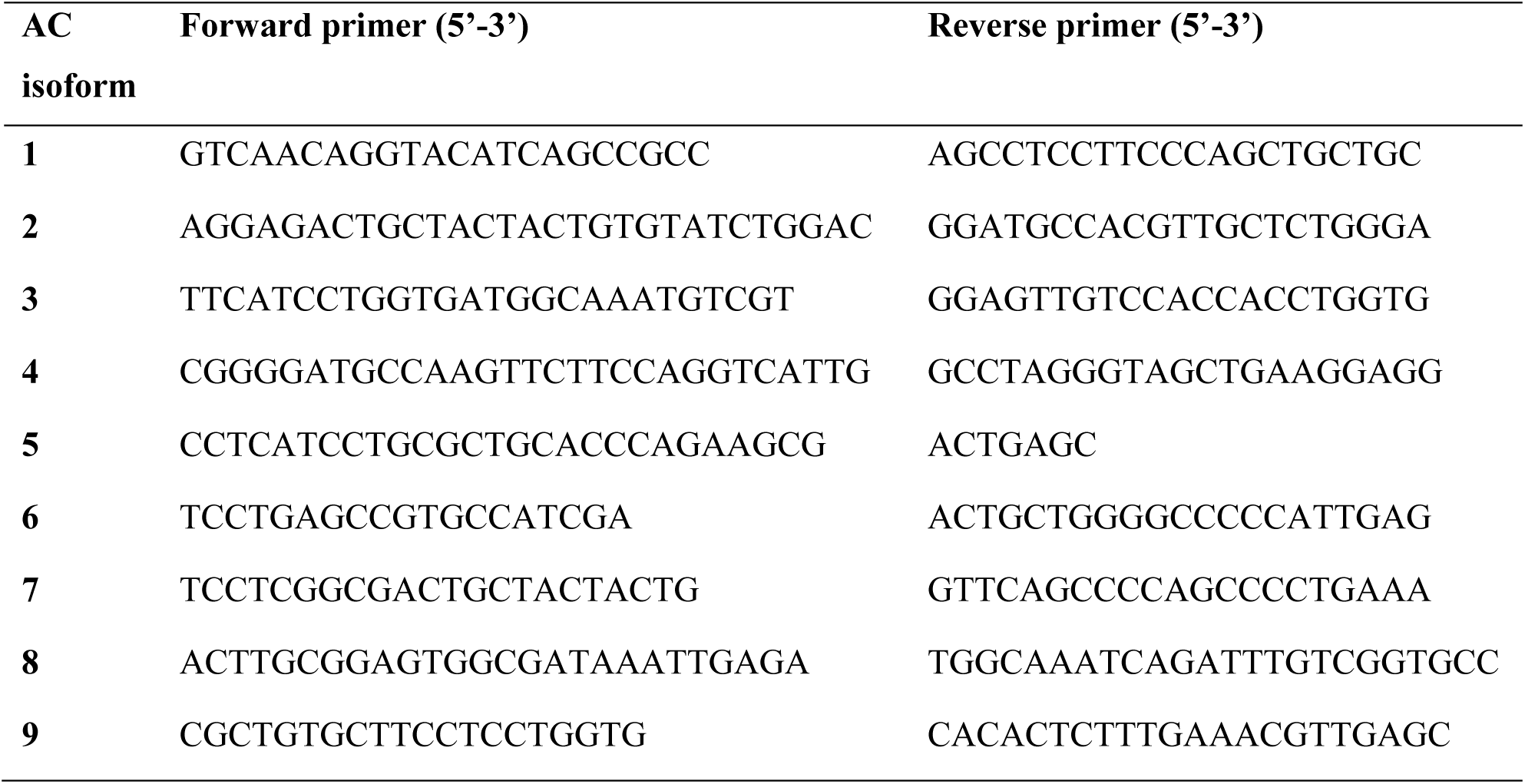

### Adenylyl cyclase assay

In a volume of 10 μl, AC activities were measured using 1 mM ATP, 2 mM MgCl_2_, 3 mM creatine phosphate, 60 μg/mL creatine kinase, and 50 mM MOPS pH 7.5. The cAMP assay kit from Cisbio (Codolet, France) was used for detection according to the supplier’s instructions. For each assay a cAMP standard curve was established. EC_50_ and IC_50_ values were calculated by GraphPad Prism version 8.4.3 for Windows, GraphPad Software, San Diego, California USA, www.graphpad.com.

### cAMP accumulation assay

HEK293 cells stably expressing mAC isoforms 3, 5, and mAC5(membr)_mAC3(cat) were plated at 2500-10000 cells/well into 384 well plates. Cells were treated with varying concentrations of lipids and incubated for 10 min at 37°C and 5% CO_2_. 2.5-10 μM isoproterenol was added to stimulate cAMP formation and cells were incubated for 5 min. HEK293-AC5-3 was assayed in the presence of the phosphodiesterase inhibitor 0.5 mM isobutyl-methyl-xanthine. Addition of Cisbio HTRF detection reagents stopped the reaction and cAMP levels were determined.

### Data handling and analysis

Experimental results were evaluated using GraphPad Prism version 8.4.3. Assays were conducted with a minimum of two technical replicates from at least two independent assays, as specified in the figure legends. Average values from duplicate or triplicate experiments were designated as single points and data are expressed as means ± S.E.M. Graphs were generated by GraphPad and assembled in PowerPoint.

## Acknowledgments

We are indebted to Prof. Andrei Lupas, Max-Planck-Institute of Biology, Tübingen, for support, advice and critique. We to thank N. Grzegorzek, Organic Chemistry, for GC/MS measurements. Funding was from the Deutsche Forschungsgemeinschaft (Schu275/45-1) and from institutional funds from the Max-Planck-Gesellschaft;

## Author contributions

Experimental realization: ML, SE and AS. HPLC, GC/MS, NMR, SE and HG; Connectase assay, AF; Concept, data evaluation and writing JES.

Conflict of interest. The authors declare no conflict of interest with the contents of this article.

Data and Materials availability: All data are available in the paper and the two appendices.

